# Multivariate prediction of cognitive performance from the sleep electroencephalogram

**DOI:** 10.1101/2023.02.28.530401

**Authors:** Péter P. Ujma, Róbert Bódizs, Martin Dresler, Péter Simor, Shaun Purcell, Katie L. Stone, Kristine Yaffe, Susan Redline

## Abstract

Human cognitive performance is a key function whose biological foundations have been partially revealed by genetic and brain imaging studies. The sleep electroencephalogram (EEG) is tightly linked to structural and functional features of the central nervous system and serves as another promising biomarker. We used data from MrOS, a large cohort of older men and cross-validated regularized regression to link sleep EEG features to cognitive performance in cross-sectional analyses. In independent validation samples 2.5-10% of variance in cognitive performance can be accounted for by sleep EEG features, depending on the covariates used. Demographic characteristics accounted for more covariance between sleep EEG and cognition than health variables, and consequently reduced this association by a greater degree, but even with the strictest covariate sets a statistically significant association was present. Sigma power in NREM and beta power in REM sleep were associated with better cognitive performance, while theta power in REM sleep was associated with worse performance, with no substantial effect of coherence and other sleep EEG metrics. Our findings show that cognitive performance is associated with the sleep EEG (r=0.283), with the strongest effect ascribed to spindle-frequency activity. This association becomes weaker after adjusting for demographic (r=0.186) and health variables (r=0.155), but its resilience to covariate inclusion suggest that it also partially reflects trait-like differences in cognitive ability.

## Introduction

Cognitive performance in humans is a fundamental neuropsychological function which predicts both sociological outcomes ^1–3^ and the development or progression of disease ^4^. Human cognitive performance varies, among others, as a consequence of genetic factors ^5^, long-acting environmental influences like schooling or toxin exposure ^6, 7^, proximal environmental factors such as stress or sleep deprivation ^8–10^, as well as various somatic and psychiatric illnesses ^11–13^.

Finding the biological foundations of individual differences in cognitive performance has been a mainstay of neuroscience research for the past decades ^13, 14^. Early studies of human cognitive biomarkers have typically been conducted in small samples analyzed with non-standardized methods, a problem generally present in the psychological ^15, 16^, and neuroscience literature^17, 18^.When biomarkers of human cognition have low effect size – that is, individual differences in cognitive performance arise due to the summed effects of many small biological differences – then the signal-noise ratio in the detected associations is poor, leading to many false positive findings which do not replicate while the true associations may remain undetected ^19, 20^. Recently, however, large-scale studies have explored some biomarkers of human cognition, especially genetic ^21, 22^ and magnetic resonance imaging (MRI) related ^23–27^ features. These studies have revealed some biological characteristics linked to various aspects of human cognitive performance, however, as they account for a small fraction of the variance, they currently do not provide a full mechanistic description about the origin of individual differences in cognitive performance.

The sleep electroencephalogram (EEG) is another important biomarker of cognitive performance. This is for two main reasons. The first reason that while the sleep EEG changes substantially over the course of the human lifespan ^28–31^, sleep EEG measures obtained from the same individual on different nights within a reasonably short time period are highly similar ^32–35^ even if the night or the preceding day are perturbed ^36^, while they exhibit substantial inter-individual variability. In other words, sleep EEG features are trait-like, which renders them strong potential candidate biomarkers of other temporally stable traits. The second reason relates to the biological significance of the EEG signal. The sleep EEG, when recorded from the scalp, reflects the joint activity of relatively large neuronal assemblies in the underlying brain tissue with excellent temporal (although limited spatial) precision. Thus, the sleep EEG can provide information about both the structural ^37–40^ and functional ^41, 42^ features of the central nervous system which are potentially not available for other imaging modalities. Notably, certain oscillations detectable from the sleep EEG – most importantly, slow waves and sleep spindles – have physiologically clearly described generating mechanisms and, in part, functions^42, 43^. If these oscillations are associated with cognitive performance or another human characteristic, then this provides mechanistic information about the biological foundations of this trait and may highlight intervention targets if the trait is clinically relevant. The trait-like nature and intimate link to both structural and functional features of the central nervous system render the sleep EEG a highly promising biomarker of other individually stable human characteristics linked to the central nervous system, such as cognitive performance.

Despite its potential, the sleep EEG is somewhat underutilized in the search for cognitive biomarkers, although this is changing with the advent of large, freely available sleep EEG cohorts ^44^. Some literature, however, has clearly linked the sleep EEG to cognitive performance. Notably, the sleep EEG can be linked to cognitive performance for at least two different reasons, both of which are potentially significant, although more so for two different fields of scientific inquiry and with different applications.

First, both sleep and cognitive performance changes as a function of age ^41, 45^, as well as in association with various common health conditions ^12, 46^. Thus, any association between sleep features and cognitive performance can arise due to age or poor health serving as either a common cause or a mediating factor in both. For example, aging may lead to both a reduction of slow wave sleep and worse cognitive performance (common cause), or obesity may lead to poor sleep and this in turn may lead to poor cognitive performance (mediation). These associations are likely to arise in geriatric sleep cohorts where mean participant age is high, its variability is considerable and participants frequently suffer from age-related ailments. A recent landmark study ^47^ has explored the association between the sleep EEG and cognitive performance considering these issues. The study found that numerous features of the sleep EEG, such as features of sleep spindles and slow waves, were associated with cognitive performance, even after correcting for chronological age and health-related covariates. It also reported that sleep EEG features associated with higher age are also generally associated with worse cognitive performance, even after correcting for age.

Second, a line of research has linked sleep in general and the sleep EEG in particular to psychometric intelligence ^48–52^, generally in healthy young participants where comorbidities were not likely an issue. A meta-analysis linked the amplitude of sleep spindles to higher scores on IQ tests ^53^. Individual studies found that spectral features were ^54, 55^, but coherence ^56^ was not associated with IQ test performance in healthy participants. These studies also highlighted the role of spindle-frequency oscillations in cognitive functioning.

Our goal in the current study was to utilize a data-driven, cross-validated approach in a large sleep EEG cohort to link features of the sleep EEG to general cognitive performance, defined as the first principal component of multiple cognitive test scores, while taking into account multiple sets of covariates which may account for some of this association.

## Methods

### Electroencephalography recordings

For our principal exploratory analyses, we used data from the MrOS Sleep Study. MrOS Sleep is an ancillary study of the parent Osteoporotic Fractures in Men Study (MrOS) ^57, 58^. Between 2000 and 2002, 5,994 community-dwelling men 65 years or older were enrolled at 6 clinical centers in a baseline examination (mean age in current sample: 73.06 years, SD=5.55 years). Between December 2003 and March 2005, 3,135 of these participants were recruited to the Sleep Study when they underwent full unattended polysomnography and 3 to 5-day actigraphy studies ^59, 60^. In these studies, EEG was recorded from C3 and C4 with a sampling frequency of 256 Hz, a high-pass hardware filter of 0.15 Hz. Both channels were recorded with gold cup electrodes, originally referenced to Fpz, and re-referenced to the contralateral mastoids. All recordings were visually scored by experts (see ^47^ for further recording details). Artifacts were automatically rejected. Artifact rejection was performed on a 4-second basis. The three Hjorth parameters were calculated for all 4-second epochs and those deviating from the within-participant mean of the given vigilance state (NREM or REM) by at least 2 standard deviations were rejected as artifactual ^61^. The selection of participants for the current study is illustrated on Figure 1.

**Figure 1.**
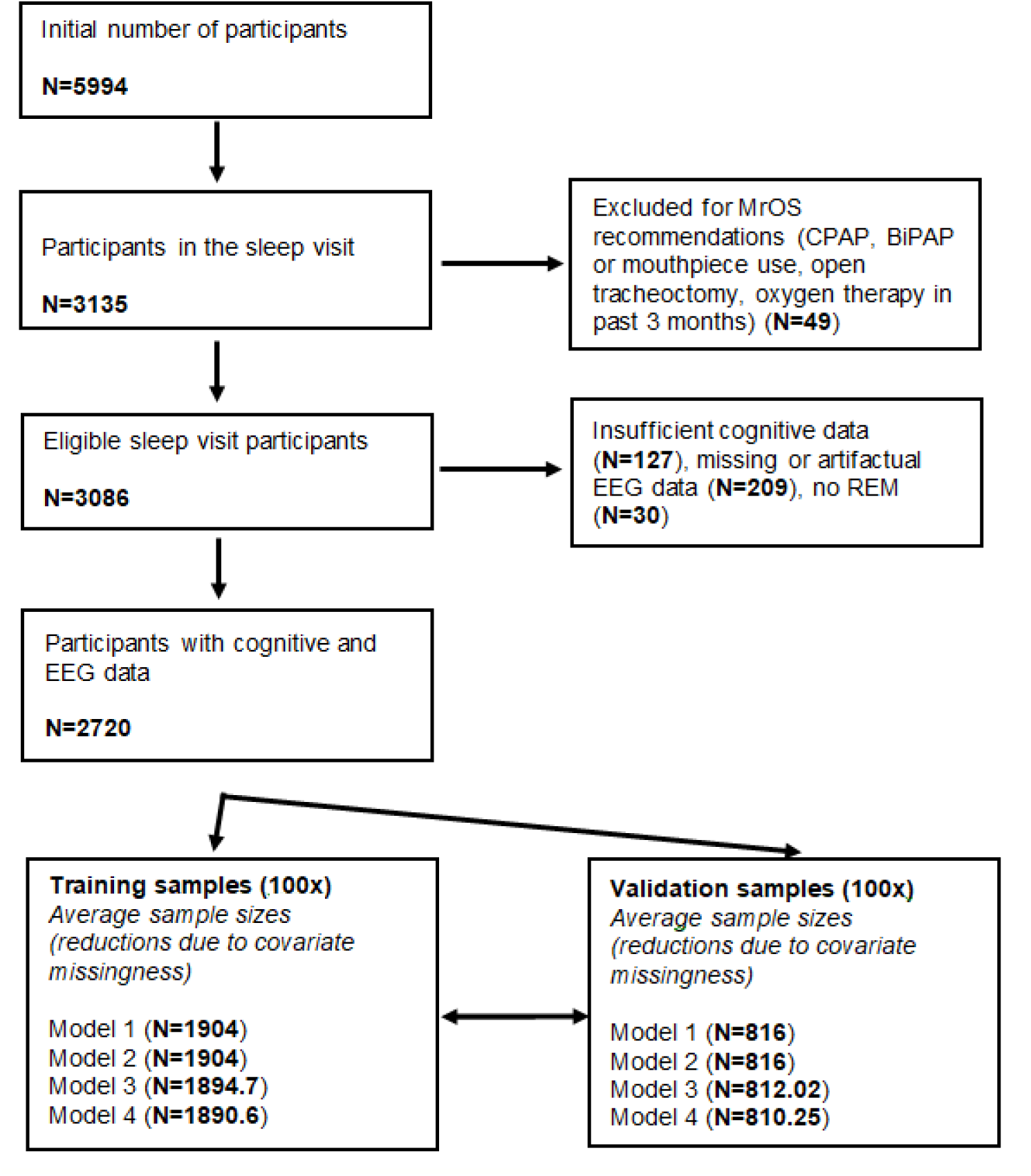
Sample size flowchart. Fractional sample sizes in the training and validation sample sets reflect the fact that across the 100 random splits some variation in sample size was observed.

### EEG feature extraction

From the EEG data, we extracted a set of features as intended predictors of general mental functioning. This set of predictors was selected as plausible correlates of cognitive performance based on previous literature (see Introduction).

1. Power spectral density (PSD), 0-48 Hz with 0.25 Hz bin resolution, separately for C3 and C4 and in REM and NREM sleep. All PSD estimates were log10 transformed and relativized by subtracting the mean of all PSD values (across bins, within each participant, channel and sleep state) from all individual PSD values.
2. PSD laterality: C3-C4 PSD difference in both NREM and REM, calculated from the averaged, relativized data.
3. NREM-REM PSD difference on the mean of both channels, calculated from the averaged, relativized data.
4. Weighted phase lag index (wPLI), 0-48 Hz with 0.25 Hz bin resolution, calculated between C3 and C4 in REM and NREM sleep. wPLI ^62^ is a measure of signal synchronization in two sources which is designed to penalize zero phase lags to reduce the effects of spurious signal similarity due to volume conduction.
5. Hjorth parameters ^63^ activity, mobility and complexity on both channels, in REM and NREM separately. Hjorth parameters are simple measures describing the stationarity of a signal, and estimate the total power, the mean frequency and the bandwidth of the signal, respectively.
6. The Modulation Index ^64^ between delta (0.5-4 Hz) phase and sigma (10-16 Hz) power on both channels, in REM and NREM separately. Modulation Index quantifies the degree to which a higher-frequency signal is modulated by a lower-frequency signal. In this case, this measure was intended to estimate the degree to which sleep spindles are coupled by slow waves ^65^.
7. Linear and quadratic overnight trends for all previous predictors. These were estimated by regressing time since recording onset at the epoch start on the predictor values calculated from each epoch. For this we estimated magnitude-squared coherence for each epoch, using the mscohere() MATLAB function and splitting each epoch into eight overlapping windows to gain a within-epoch estimate of coherency. In order to simplify analyses, for PSD and wPLI we calculated the average delta (0.1-4 Hz), theta (4-8 Hz), alpha (8-10 Hz), low sigma (10-13 Hz), high sigma (13-16 Hz), beta (16-25 Hz) and gamma (25-48 Hz) power in each epoch instead of regressing time on power in each individual frequency bin. Frequency bins on the borders between frequency bands were included in the calculation of the higher frequency band.

All features (except wPLI) were calculated for each 4-second epochs in the signal with 50% overlap and then averaged across windows to yield a single value in each participant. For wPLI, the imaginary part of the cross-spectrum was calculated for each window in this manner, and then averaged by using real components as weights according to the formula provided by Vinck et al. For PSD and wPLI, Hamming windowing was used.

For a less fine-grained analysis of EEG spectral components, we averaged PSD estimates within the following ranges to obtain band power: delta (0.1-4 Hz), theta (4-8 Hz), alpha (8-10 Hz), low sigma (10-13 Hz), high sigma (13-16 Hz), beta (16-25 Hz), gamma (25-48 Hz). Power at band boundaries was always assigned to the higher-frequency band.

### EEG spectral parametrization

Spectral components of the sleep EEG do not necessarily reflect actual oscillations ^66, 67^. Much of the variance in spectrum of the sleep EEG can be modelled with just two parameters, a spectral intercept and a slope coefficient describing the exponent of and 1/f power law function (aperiodic components). Oscillations cause a deviation from this deterministic pattern (periodic components). We used FOOOF ^66^ to decompose absolute spectra into periodic and aperiodic components, estimating the power law function in the full (0.25-48 Hz) range. We allowed periodic components (spectral peaks) with a width of 0.5-6 Hz and a minimum peak height of 2 standard deviations above the aperiodic spectrum. We discarded participants for whom periodic and aperiodic components accounted for less than 95% of the variance in the power spectrum (N=81). Spectral parametrization was performed separately on both EEG channels and in NREM and REM sleep. Based on the zero-order correlations between cognitive performance and EEG power (see Results) we searched for peaks in the REM theta, REM beta and NREM sigma ranges. The bandwidth, frequency and power (height above the aperiodic spectrum) of these peaks was saved. If a participant did not have a detected peak in these ranges, the value of the spectral peak was set to 0 and the value of the bandwidth and frequency were set to the sample average. If a participant had multiple peaks in these frequency ranges, we retained the one closest in frequency to the maximum of the zero-order correlation between power spectral density and cognitive performance (6.5 Hz in REM theta, 23 Hz in REM beta and 14 Hz in NREM sigma).

### Cognitive data

Concurrent with the sleep study, participants filled out three cognitive tests: the Modified Mini-Mental State Test (3MS), Trails B, and Digit Vigilance (DV). From these tests, we considered the following variables: 3MS total score, Trails B completion time, DV completion time, and DV omission errors (false negatives). 3MS total scores were square root-transformed to improve normality and their inverse was taken to ensure that in all tests higher scores mean worse performance. The other scores were used without transformation.

In this sample, raw cognitive test scores may have been strongly affected by factors other than general mental functioning, most notably age and health. Consequently, we regressed out a set of covariates from the raw scores. Because (with the exception of age) the route of causation between the confounding variables and test scores is unclear, we explored four models with four, increasingly extensive sets of covariates:

- Model 1: no covariates
- Model 2: regressing out technical/demographic variables (recording site, age including quadratic, cubic and fourth-order effects, and race/ethnicity)
- Model 3: regressing out technical/demographic variables, plus health (medication use, systolic and diastolic blood pressure, caffeine, alcohol and cigarette consumption before sleeping, comorbidities listed at the baseline visit [arthritis/gout, cancer, cataracts, congestive heart failure, diabetes, glaucoma, kidney stones, osteoporosis, Parkinson’s, prostatitis, stroke], comorbidities listed at the Sleep Study [angina pectoris, peripheral, cerebral or coronary disease, arterial fibrillation, heart rate problems, sleep disorders]). At both the baseline and sleep visits, participants answered a questionnaire about comorbidities with the following formula: “Have you ever had (disease name)?”, except angina pectoris, which was measured with the Rose Angina Questionnaire. In order to reduce missing data, we considered participants not providing information about a comorbidity to not have that condition. We also used the use of 49 common medications (based on a physician’s review of the participant’s common medications presented during a personal visit) as covariates (see Supplementary text for a detailed list).
- Model 4: regressing out demographic/technical variables, physical health and quality of life, including mental health and sleep symptoms (SF12 Modified Physical/Mental Summary Scale score, Geriatric Depression Scale score, Goldberg Anxiety and Depression Scale scores, Epworth Sleepiness Scale scores, Pittsburgh Sleep Quality Inventory total score, Functional Outcomes of Sleep Questionnaire total score). For simplicity, we refer to the covariates only included in Model 4 as ‘quality of life’.

In all models, we also regressed out the effect of confounders from the EEG predictors.

### Statistical analysis

In our main analyses, the principal question was: to what extent can cognitive performance be predicted from sleep EEG biomarkers? In order to answer this question, we used sleep EEG features as independent variables and cognitive performance as the dependent variable in regularized regression models ^68, 69^. Regularized regression is an iterative learning algorithm which minimizes the following function:

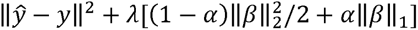

In plain words, regularized regression performs ordinary linear regression, but it also assigns a penalty to the prediction error, which increases as a function of 1) more predictors with non-zero beta coefficients in the model 2) an iteratively changing λ penalty parameter. Regularized regression can use L1 (LASSO regression) or L2 (ridge regression) regularization, or a combination of the two (elastic net regression. The combination of L1 and L2 regularization can be tuned with a parameter α. α=0 yields ridge regression and α=1 yields LASSO regression, while interim values yield elastic net regularization. Due to regularization, all of these procedures are able to handle more predictors than there are cases, while robust results are ensured and overtraining is protected against by cross-validation.

We randomly split all MrOS participants into a training (70%) and a validation (30%) sample. Regularized regression models were fitted with 10-fold cross-validation in the training sample. We split the training sample into 10 random subsamples and iteratively fitted the regression model using a range of λ values (penalty parameters) in nine of them, with the tenth serving as internal validation control to assess performance. This was repeated in all combinations of subsamples.. The effect size of interest (henceforth referred to as validity) was the correlation between predicted and actual cognitive functioning in the validation sample. We performed this analysis 100 times to explore the effect of randomly assorting participants into training and validation samples. We repeated this procedure for dependent and independent variables after regressing out the effects of each covariate set (Model 1-4), and across a range of α values (0-1 with increments of 0.1) that switch between ridge (α=0), elastic net (0<α<1) and LASSO (α=1) regressions, yielding 44 model specifications (covariate sets and α) and 100 models with random subsamples for each specification.

Regularized regression was performed using the cvglmnet() MATLAB function, based on the glmnet() package ^70^. Due to missingness of data, the number of participants slightly varied in the validation samples, but it was on average N=816 in Model 1 and 2, N=812 in Model 3 and N=810.25 in Model 4 (minima: N=793, N=793, N=788, N=786, respectively).

### Data and code availability

All PSG data are freely available via the National Sleep Research Resource (http://sleepdata.org). Model results and code used for analyses are available on Zenodo at 10.5281/zenodo.7684266.

## Results

### Covariate effects on cognition and sleep

Potential covariates accounted for up to 20% of the variance in cognitive scores, with the least in Digit Vigilance errors and the most in Trails B completion time (Figure 2). About two thirds of this variance in 3MS, Digit Vigilance completion time and Trails B completion time and virtually all of this variance in Digit Vigilance errors was accounted for by demographic covariates alone. Supplementary figures 1-4 provide detailed data about the association between covariates and power spectral density. Supplementary figure 5 illustrates the relationship between covariates and cognitive test scores.

**Figure 2.**
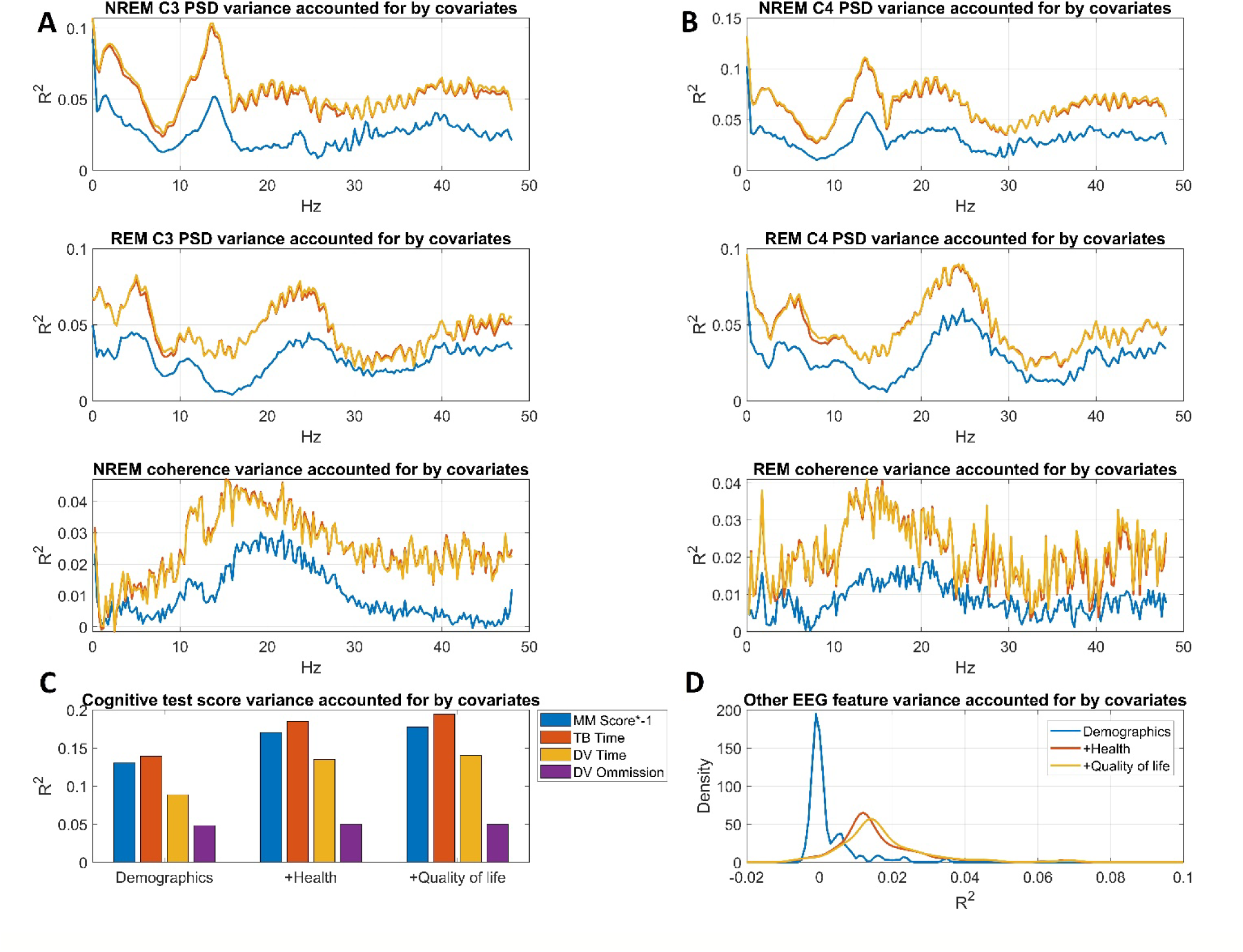
Variance accounted for by the three covariate sets (Model 2: demographic covariates added, Model 3: health covariates added, Model 4: quality of life covariates added). Panel A shows power spectral density variance accounted for as a function of frequency. Panel B shows wPLI variance accounted for as a function of frequency. Panel C shows variance accounted for in raw cognitive test scores. Panel D shows the distribution of R^2^ values across the 190 other EEG features.

In sleep measures, potential covariates accounted for up to 10% of the total variance, with about half attributable to demographics. Covariates accounted for the most variance in NREM delta, NREM sigma, REM theta and REM beta power and in wPLI values from a broad frequency range encompassing the sigma and beta bands. Notably, health-related covariates affected sleep measures in a very similar manner to age, and quality of life accounted for very little additional variance, although it encompassed explicit questionnaire-based measures of sleep.

### Principal component analysis of test scores

Cognitive test scores were all positively correlated, with or without regressing out covariates (Supplementary table S1). In all models, we performed principal component analysis on the unstandardized residuals of cognitive scores. In all models, a single principal component with eigenvalue>1 emerged. This first principal component accounted for 42.5-46.3% of the variance, with values decreasing somewhat with the inclusion of more confounders. In each model, we extracted principal component scores on this first unrotated principal component as the measure of general cognitive performance. Measures from the 4 models were highly correlated (Supplementary table S2, r=0.84-0.99), in line with the observation that the confounders only accounted for a modest amount of variance in test scores.

### Correlations between the sleep EEG and cognition

In our initial analysis, we calculated zero-order correlations between general cognitive functioning and sleep measures in the entire MrOS sample. Results were in line with previous reports ^47, 54^. In NREM sleep, higher relative power in the broad sigma (∼8 Hz-20 Hz) range was associated with better cognitive performance, with a clear peak in the fast spindle range around 14 Hz. High-frequency power (>30 Hz) was associated with lower cognitive performance. In REM sleep, power in the beta range (∼18-32 Hz) was associated with better cognitive performance, while higher power in the theta (∼5-7 Hz) range was associated with lower performance. This pattern of results was consistent across the four models with different covariate sets, although effect sizes were reduced in more heavily corrected models and only reached r>0.1 for the NREM sigma association in Model 4 (demographic, health and quality of life covariates).

We found no consistent correlations between cognitive function and wPLI values or other EEG features. Figure 3 illustrates bivariate correlations between cognitive performance and sleep EEG measures.

**Figure 3.**
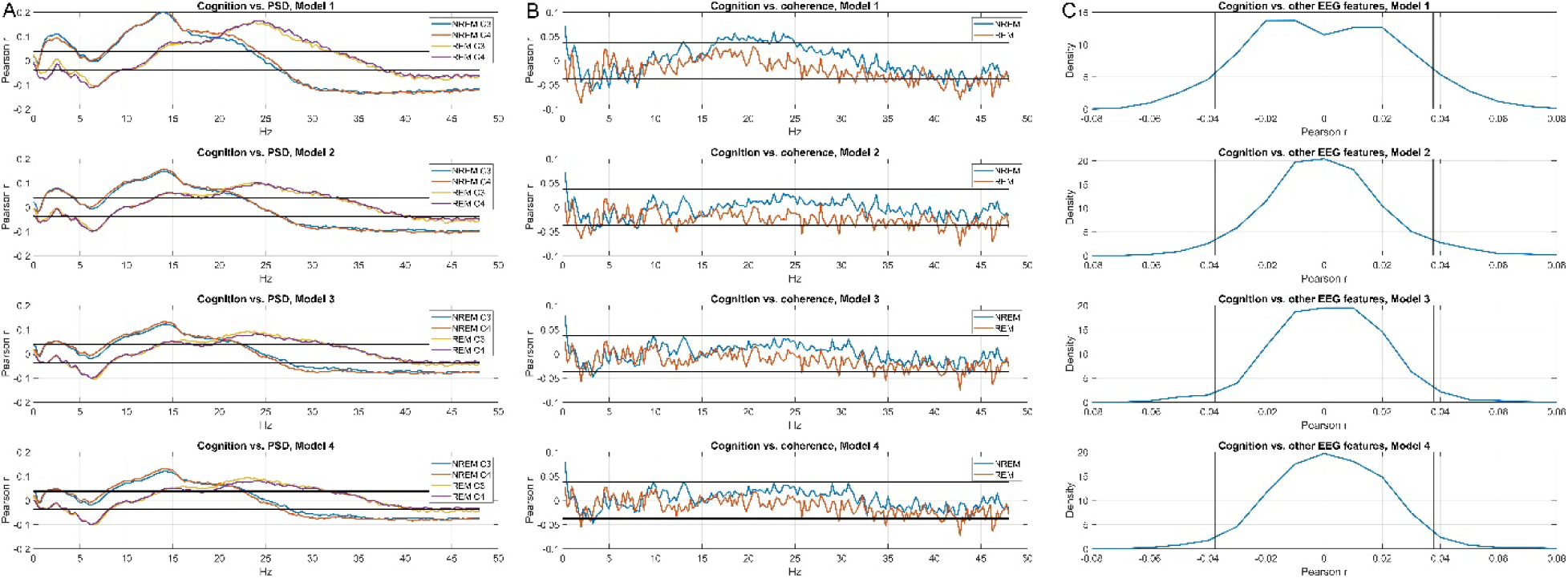
Correlations between general cognitive performance, power spectral density (Panel A), wPLI-based functional connectivity (Panel B) and other EEG features (Panel C). Correlations are shown after increasingly strict covariate sets: no covariates (Model 1), demographic covariates added (Model 2), health covariates added (Model 3), quality of life covariates added (Model 4). For PSD and wPLI, correlations are shown as a function of frequency. For other EEG features, only the distribution of correlation is shown for simplicity, as no correlation is significant after adjusting for multiple comparisons. On all plots, black lines indicate the critical correlation coefficient.

### The sleep EEG predicts cognitive performance

In our initial models, we used ridge regression (α=0) to predict general cognitive functioning. We used 70% of MrOS participants as training and 30% as validation, repeating this process 100 times to get an estimate of the variation in model performance due to random sampling of the training and validation samples. Figure 4 provides a detailed illustration of prediction performance. Across the 100 random samples, the mean validity (out-of-sample correlation between predicted and actual cognitive performance) amounted to 0.283 (SD=0.026, range 0.219- 0.359) in Model 1 (no covariates), 0.186 (SD=0.027, range 0.084-0.246) in Model 2 (demographic covariates added), 0.155 (SD=0.026, range 0.077-0.211) for Model 3 (health covariates added), and 0.152 (SD=0.026, range 0.067-0.212) for Model 4 (quality of life covariates added) (Figure 4, Panel A). Empirical p-values can be considered to be 0 as no model had non-positive validity, but with 100 model runs we had a limited resolution of possible empirical p-values. A more conservative, semi-parametric p-value was calculated by considering the standard deviation of validity across models to be an empirical standard error. Dividing the mean validity by this number to obtain a z-statistic and converting it into a p-value yields p-values of <10^−13^, 8*10^−12^, 5*10^−9^, and 7*10^−9^ for Models 1-4, respectively. We note that individual models were usually all highly statistically significant, as across 400 model runs only a single non-significant p-value (p=0.057) was observed and only 16 p-values exceeded 0.001.

**Figure 4.**
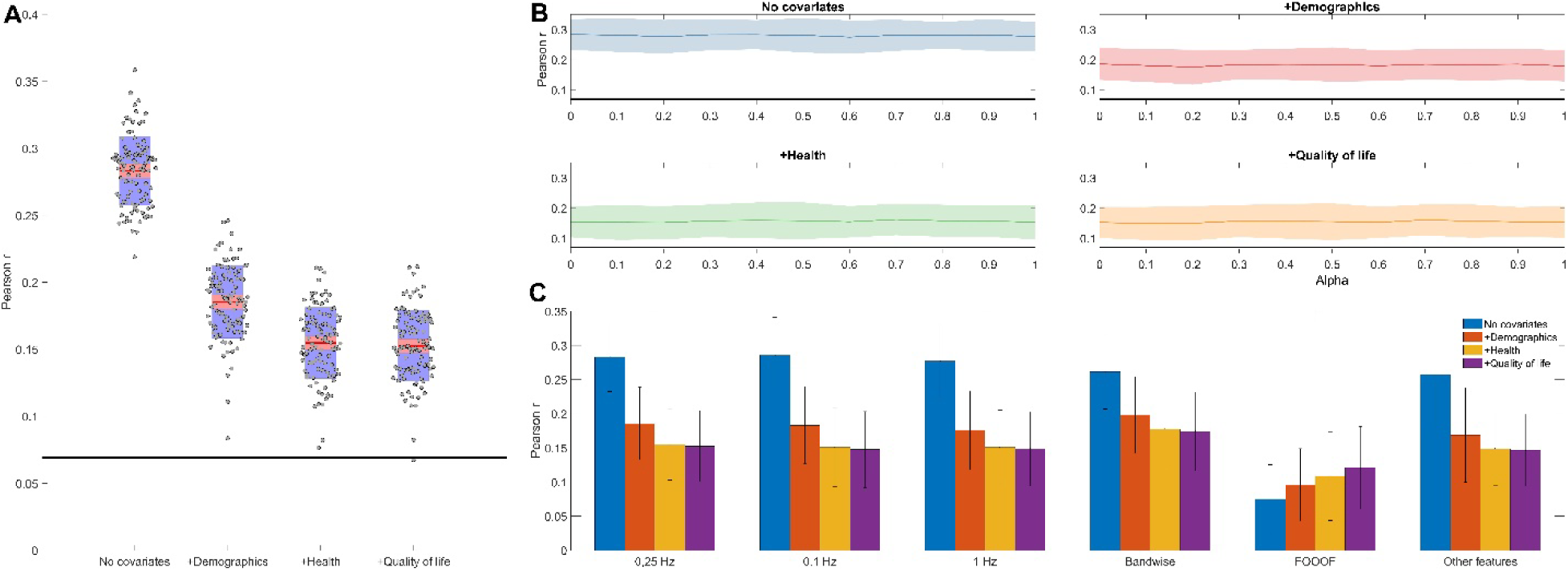
Panel A. EEG-based predictive validity (out-of-sample correlation between predicted and actual cognitive performance) for four increasingly strict covariate sets. Model 1 contains no covariates, Model 2 controls for demographic covariates, Model 3 adds controls for health covariates, while Model 4 adds controls for quality of life covariates. Red lines show the average performance across 100 model runs with random training-validation splits. Grey dots show individual model runs. Red shading illustrates the standard error of the mean, while blue shading illustrates the standard deviation across model runs. A thick black line illustrates the critical correlation coefficient at the mean sample size of validation samples. Panel B: out-of-sample correlations between predicted and actual cognitive functioning as a function of regularization type and covariate choice. Regularization type is iteratively changed between alpha=0 (ridge regression), 0<alpha<1 (elastic net) or alpha=1 (LASSO). A thick black line indicates the critical correlation coefficient at the mean sample size of the validation samples. Shadings indicate empirical standard errors (1.96 standard deviations of the out-of-sample correlations calculated from 100 model runs). Panel C: model validity with different predictors sets: binwise PSD with three different spectral resolutions (0.1, 0.25 and 1 Hz), bandwise PSD, spectral parameters (intercept, slope, bandwidth, frequency and power of REM theta/beta and NREM sigma peaks) derived by FOOOF, and 0.25 Hz binwise PSD with the other EEG features (see Methods) added.

### Alpha tuning has no effect on model performance

We found no evidence that models using α>0 (elastic net or LASSO) performed better than α=0 (ridge regression). Models using such values produced validities significantly different from ridge regression in only three cases out of the 40 comparisons: for Model 1 α=0.6 produced significantly lower validity than α=0 (β=-0.008, p=0.024); for Model 2 α=0.2 produced significantly lower validity than α=0 (β=-0.01, p=0.006); while for Model 4, α=0.7 produced significantly higher validity than α=0 (β=0.009, p=0.028). As these deviations were rare, small in magnitude and did not fit into a theoretically expected or empirically observed pattern, we considered them to be likely spurious and proceeded with the computationally simpler ridge regression as our preferred model. Figure 4, Panel B summarizes the performance of various regularized regression models.

### EEG features other than PSD lack predictive value

In further steps, we explored whether the predictive performance of the sleep EEG changes by adding further features or changing their resolution. First, we compared models based only on PSD data with models that also incorporated wPLI, phase-amplitude coupling, Hjorth parameters as well as linear and quadratic overnight trends as predictors. We found that these more complex models actually statistically significantly underperformed relative to PSD-only models in Model 1 (β=-0.026, p=3*10^−7^) and Model 2 (β=-0.017, p=2*10^−4^), while there was a statistically non-significant trend for better performance in PSD-only models in Model 3 (β=0.006, p=.0126) and Model 4 (β=0.006, p=0.122) (Figure 4, Panel C). That is, PSD remained the best predictor of cognitive performance with no meaningful additional variance accounted for by other predictors.

### Spectral resolution does not affect predictive validity

In the next step, we run models based on PSD data using two additional levels of PSD resolution (1 Hz and 0.1 Hz using zero-padding). Models based on the more sparse PSD (1 Hz resolution) tended to yield slightly lower out-of-sample correlations (β=-0.01--0.004), but this difference only reached significance in Model 2 (β=0.01, p=0.016). Models with the fine-grained PSD (0.1 Hz resolution) did not produce even a consistent trend for higher validity values (β=- 0.005-0.002 for the four models, pmin=0.22) (Figure 4, Panel C). Thus, 0.25 Hz remained our preferred resolution for binwise analysis.

### Band power has comparable predictive validity to binwise power

We attempted using band power in seven frequency bands (see Methods for details) as predictors of general cognitive functioning. It is of note that from these models we dropped not only the fine-grained power estimates, but also PSD laterality and REM-NREM PSD differences, using just 28 predictors (power in seven frequency bands over two channels in NREM and REM) in the regularized regression model. We found that these models significantly underperformed in Model 1 (β=-0.02, p=10^−8^), but actually outperformed binwise models in Model 2 (β=0.01, p=0.002), Model 3 (β=0.02, p=10^−8^) and Model 4 (β=0.02, p=10^−8^). The mean out-of-sample correlations across the 100 runs yielded by band power models were 0.262 (SD=0.028) for Model 1, 0.198 (SD=0.029) for Model 2, 0.178 (SD=0.029) for Model 3 and 0.174 (SD=0.029) for Model 4 (Figure 4, Panel C).

### Spectral parametrization

Spectral components of the sleep EEG do not only reflect oscillatory components, but also background activity and sinusoidal components introduced by Fourier analysis do approximate non-sinusoidal oscillations in the actual signal ^66, 67^. Therefore, as an alternative analytical strategy, we attempted to decompose spectra into aperiodic (an intercept and an exponent to describe non-oscillatory activity in a simple power law function) and periodic (oscillations exceeding the trend of the power law function) components and use these as predictors of cognitive functioning. Spectral parametrization was performed using FOOOF ^66^. In each participant, on each channel and in NREM and REM separately we calculated spectral intercepts, spectral slopes, as well as the bandwidth, frequency and power of REM theta, REM beta and NREM sigma peaks.

Results confirmed the findings related to PSD analyses. In univariate analyses, across all four covariate sets, better cognitive performance was significantly associated (after correction for multiple comparisons) with a higher spectral intercept, a steeper spectral slope, higher power in the NREM sigma and lower power in the REM theta ranges, and higher REM beta power on C3. The correlation between REM beta power on C4 was only found in Model 1 (no covariates) and Model 2 (demographic covariates). No other spectral parameter was consistently associated with cognitive performance, but a trend emerged between a higher-frequency REM beta peak and better cognitive performance (Figure 5, Panel A). These findings were replicated for peak missingness. Lacking NREM sigma or REM beta peaks was significantly associated with lower cognitive performance, while lacking REM theta peaks showed a trend level association with higher cognitive performance (Figure 5, Panel B), mirroring both PSD-based analyses and bivariate correlations with spectral parameters.

**Figure 5.**
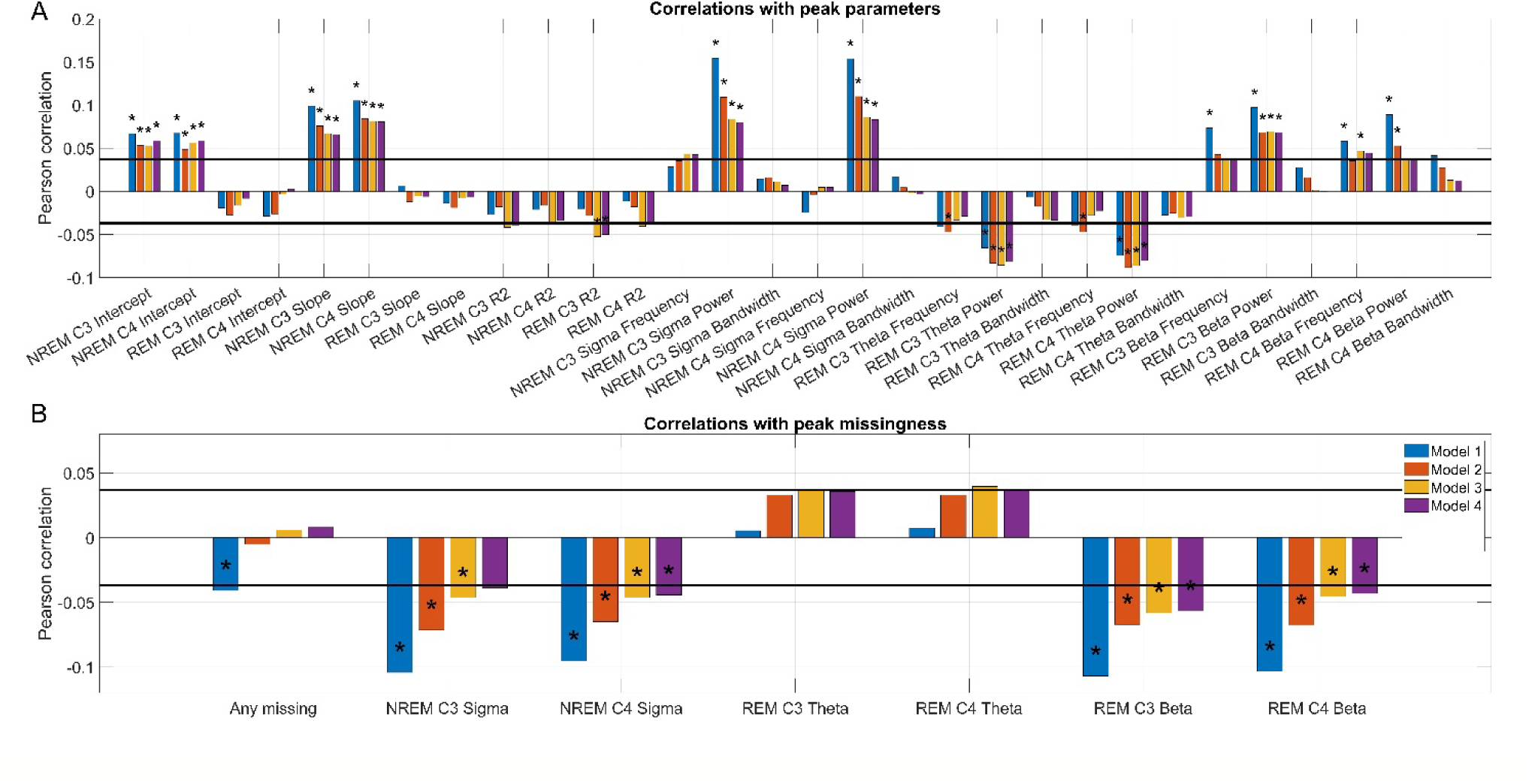
Panel A: Bivariate correlations between spectral parameters (intercept, slope, variance accounted for, and the frequency, power and bandwidth of NREM sigma, REM theta and REM beta peaks) and cognitive performance, residualized for four covariate sets. Dual horizontal lines illustrate the critical significance level assuming zero missingness. Asterisks mark correlations which remain significant after controlling for multiple comparisons. Panel B: Point-biserial correlations between the missingness of spectral peaks (NREM sigma, REM theta and REM beta peaks) and cognitive performance, residualized for four covariate sets. Dual horizontal lines illustrate the critical significance level assuming zero missingness. Asterisks mark correlations which remain significant after controlling for multiple comparisons.

As before, we trained regularized regression models to predict cognitive performance from EEG spectrum parameters, while controlling for the potential confounders specified in Models 1-4. Multivariate models based on these spectral parameters underperformed relative to models based on the 0.25 Hz PSD (Figure 4, Panel D). Out-of-sample correlations [standard deviations] with cognitive performance were: 0.075 [0.026], 0.096 [0.027], 0.109 [0.033] and 0.121 [0.031] for Models 1-4, respectively, all differences from the 0.25 Hz PSD model are significant at p<0.001.

## Discussion

Our study used a hypothesis-free, multivariate, cross-validated method to identify markers of cognitive functioning from the sleep electroencephalogram. We found that such markers exist, they mostly consist of power spectral density in the NREM sigma and REM theta and beta power, a part of their effect is mediated by observed demographic and health-related covariates and quality of life, but another part is independent from these.

Cognitive functioning is a significant human trait which predicts sociological outcomes ^3^, the development and progression and disease ^4^, but a decline in which is also the symptom of various pathological conditions ^13^. A search for biomarkers of cognitive function has been ongoing for decades. Early studies searching for cognitive function biomarkers were often underpowered and univariate, with the choice of the putative biomarker motivated by the intuition of researchers. As a result, many failed to yield replicable results, even if the original findings seemed biologically plausible ^71, 72^. The response to the failure of these studies has generally been to launch hypothesis-free, cross-validated association studies which rely on very large statistical power to precisely identify even small biological effects on the target phenotype, and sum of many small biological effects to yield a predictive score whose power is assessed in an independent sample. The most prominent hypothesis-free, cross validated studies have been genome-wide association studies (GWAS) using genetic data ^73^ and whole-brain regression using magnetic resonance imaging data ^24^. Several such studies were concerned with cognitive functioning ^25, 26^. EEG-based machine learning studies have also been published ^31, 74, 75^, but to date, ours is the first to apply a hypothesis-free, cross-validated association method to sleep EEG data to identify biomarkers of cognitive functioning.

In our models, NREM sigma, REM theta and REM beta activity clearly emerged as correlates of cognitive functioning, with out-of-sample multiple correlations of r=0.15-0.3. Regarding NREM sigma activity, our works replicates a larger body of literature ^47, 53, 76^ which found associations between various aspects of cognitive performance and sleep spindles. As sleep spindles arise from thalamocortical networks ^42^, our results point to the importance of the integrity of this system as a biological prerequisite of cognitive functioning. We previously reported ^54^ that REM beta oscillations had a positive, while REM delta-theta oscillations (albeit at a lower frequency with a maximum at 3.5 Hz) had a negative association with cognitive performance. A previous analysis of the current sample ^47^ also found that REM beta power was correlated with Digit Vigilance scores, although it did not consider a composite cognitive score as the dependent variable and it failed to find a similar association in another sleep cohort. While an invasive EEG study of humans ^77^ identified a REM theta-beta network in the anterior cingulate and the dorsolateral prefrontal cortex, likely underlying the oscillations identified in our current study, more research is needed to understand the functional properties of this system. Given the power and replication issues plaguing human neuroscience ^19, 20^, it is significant that we could replicate the observation that NREM sigma, REM theta and REM beta oscillations are correlated to human cognitive functioning, which should facilitate research into the biological origins of these oscillations.

We observed that while approximately half of the association between sleep EEG features and cognitive functioning was accounted for by measured covariates, the other half persisted despite statistical controls for a very large number of potential moderators. The largest drop in this association was seen between the first two models, by adding demographic covariates, of which we hypothesized age to be the most significant. As age is associated with both changes in cognitive performance ^45^ and in changes in the sleep EEG ^30, 31, 78^, an algorithm may find age-related sleep biomarkers which are, in turn, also related to worse cognitive performance. This expectation was confirmed by the fact that in the second step (Model 2, demographic covariates including linear and nonlinear effects of age added) validity dropped substantially to from 0.283 to 0.186. In the third step (Model 3, health-related covariates added, including comorbidities and medication) validity dropped further, but only slightly, to 0.155. Thus, while a small amount of the sleep EEG-cognition covariance was due to some participants’ comorbidities and/or medications being related to both sleep EEG patterns and cognitive performance, this was comparatively a small effect and even among participants of the same medical history we would expect these EEG markers to be related to cognitive performance. In the fourth step (Model 4, quality of life covariates added), we did not observe a substantial drop validity, which was on average 0.152. Interestingly, the covariates added at this step not only included geriatric functioning scales (SF12 and GDS) the scores of which could be strongly related to well-preserved cognitive functioning at higher ages, but also sleep quality rating scales (ESS, PSQI and FOSQ). Results from Model 4 disconfirm the hypothesis that EEG biomarkers of cognition index poor sleep which impairs next-day cognition, or age-related cognitive and physical decline which is also reflected in sleep alterations. Controlling for the previously added covariates, self-reported sleep quality and geriatric functioning hardly mediates any of the association between sleep EEG markers and cognition.

We did not observe substantial zero-order correlations between sleep EEG biomarkers other than power spectral density, and consequently we only included this measure as a predictor in our base model. Furthermore, based on experiences from brain imaging ^24^ and genetics ^71^, we expected that predictive validity will be driven by a relatively large number of sleep EEG features, each having only a weak zero-order association with cognition. Therefore, our initial models only included PSD as a predictor, but with a relatively fine (0.25 Hz) resolution.

Relating to the first expectation, in exploratory analyses we indeed found that adding wPLI, Hjorth parameters, delta-sigma coupling and overnight effects of all predictors to our models did not improve predictive accuracy. This confirms our finding that sleep EEG functional connectivity is not significantly associated with cognitive performance ^56^, but it is in contrast with some studies, generally performed in small samples which found that delta-sigma coupling (or a more explicitly measured grouping of sleep spindles by slow waves) is associated with cognitive outcomes ^79–82^. Notably, it is also in contrast with a similar analysis of the present sample ^47^, which found that the coupling of individually detected sleep spindles and slow oscillations was associated with better performance on some cognitive tests (Trails B and 3MS). In the current analyses, the NREM Modulation Index of the delta and sigma frequency ranges was only weakly and non-significantly associated with better cognitive composite scores, but with the correct sign on both C3 and C4 (r=0.012-0.013). Our current study deliberately used measures of spectral power instead of individually detected oscillations due to the methodological issues of sleep oscillation detection ^83, 84^, especially in older samples, in particular the issues of various sleep spindle detection algorithms in capturing the spindle-cognition association ^53^. It is possible that adequately parametrized oscillation detectors yield better estimates of slow oscillation-spindle coupling than delta-sigma Modulation Index, which may be associated with cognition. This finding highlights that while in the case of some biomarkers alternative measures yield highly comparable results (for example, sleep spindle amplitude, sigma power and sigma peak height in the parametrized spectrum all correlate with cognition), in other cases it may be necessary to measure biomarker in a precisely defined way to make associations detectable.

Relating to the second expectation, however, we found tha 1) even zero-order correlations between PSD and cognitive performance are substantial, often excluding r=0.1 2) increasing spectral resolution does not improve and reducing it does not impair predictive accuracy, and 3) very similar validities can be achieved by just retaining a coarse estimate of PSD in seven frequency bands as predictors. On the other hand, we also observed that 1) using parametrized spectra (slope, intercept and three spectral peaks) as predictors did reduce validity, and 2) the use of LASSO (which assumes sparsity, forcing regression coefficients to zero for all except a few predictors from correlated sets) was not preferred to ridge regression (which distributes regression weights among correlated predictors). These observations, taken together, suggest that while the associations between sleep EEG features (especially NREM sigma and REM beta power) and cognition are orders of magnitude stronger than what is usually observed in genetics and brain imaging, the set of associated features cannot be reduced to a handful of readily observable spectral peaks or one or two frequency ranges. Power in spectral components of the sleep EEG which are assigned to the ‘aperiodic’ part of the spectrum is substantially associated with cognitive performance.

Our work suffers from a number of limitations. First, as we use a cross-sectional design, we cannot clearly ascertain routes of causation, which also pertains to covariate selection. We emphasize that although Model 1-4 uses an increasingly strict set of covariates, stricter models are not necessarily theoretically preferred. This is because various comorbidities ^85, 86^, general well-being at a high age ^4^, and even less pronounced age-related changes in the sleep EEG ^87^ have been associated with premorbid cognitive functioning. Therefore, comorbidities may not be true confounders but simply the consequences of pre-existing cognitive abilities which are subsequently reflected in both cognitive test scores and sleep EEG patterns. The theoretical case is stronger for preferring Model 2 (demographic covariates) over Model 1 (no covariates), as both age ^30, 31^, and self-reported ethnicity ^61, 88, 89^ has a likely spurious association with sleep EEG patterns, and the same can be assumed for recording site. In any case, it is clear that even with a potentially overcontrolling strict covariate set cognitive functioning is related to features of the sleep EEG. Second, although our findings are robust to a large set of health-related covariates and replicate in an independent sample, it is not fully elucidated to what extend we found sleep EEG correlates of age-related cognitive decline or those of pre-existing cognitive abilities which persisted into an old age. For this limitation to be overcome, similar investigations in healthy, younger samples are necessary. Finally, we emphasize that the associations we find between EEG patterns and cognitive performance are modest and most of the variance in cognitive performance is not accounted for by patterns of the sleep EEG.

In sum, our work showed using a large dataset and a data-driven, cross validated approach that features of the sleep electroencephalogram are related to cognitive functioning in elderly participants, even after controlling for a broad set of covariates. Power in the NREM sigma, REM theta and REM beta bands is especially strongly implicated. These features of the sleep EEG exhibit zero-order correlations often exceeding r=0.1, and similar multiple correlations to brain imaging-based or genetic predictors ^26^, usually established in discovery samples orders of magnitude larger than ours.

## Acknowledgements

The Osteoporotic Fractures in Men (MrOS) Study is supported by National Institutes of Health funding. The following institutes provide support: the National Institute on Aging (NIA), the National Institute of Arthritis and Musculoskeletal and Skin Diseases (NIAMS), the National Center for Advancing Translational Sciences (NCATS), and NIH Roadmap for Medical Research under the following grant numbers: U01 AG027810, U01 AG042124, U01 AG042139, U01 AG042140, U01 AG042143, U01 AG042145, U01 AG042168, U01 AR066160, R01 AG066671, and UL1 TR002369. The National Heart, Lung, and Blood Institute (NHLBI) provides funding for the MrOS Sleep ancillary study “Outcomes of Sleep Disorders in Older Men” under the following grant numbers: R01 HL071194, R01 HL070848, R01 HL070847, R01 HL070842, R01 HL070841, R01 HL070837, R01 HL070838, and R01 HL070839. The National Heart, Lung, and Blood Institute provided funding for the ancillary MrOS Sleep Study, “Outcomes of Sleep Disorders in Older Men,” under the following grant numbers: R01 HL071194, R01 HL070848, R01 HL070847, R01 HL070842, R01 HL070841, R01 HL070837, R01 HL070838, and R01 HL070839. The National Sleep Research Resource was supported by the National Heart, Lung, and Blood Institute (R24 HL114473, 75N92019R002).

This research has been implemented with the support provided by the Ministry of Innovation and Technology of Hungary from the National Research, Development and Innovation Fund, financed under the PD-21 (project number: 138935) and TKP2021-EGA-25 funding schemes.

