## Supplementary figure for "Multivariate prediction of cognitive performance from the sleep electroencephalogram"

| *Model 1 - No covariates* |  |  |  |  |
| --- | --- | --- | --- | --- |
|  | **MM Score** | **Trails B time** | **DV Time** | **PC1** |
| **MM Score** |  |  |  | 0,390 |
| **Trails B time** | 0,465 |  |  | 0,682 |
| **DV Time** | 0,254 | 0,459 |  | 0,577 |
| **DV Errors** | 0,137 | 0,126 | 0,085 | 0,225 |
|  |  |  |  | **R2=46,25%** |
| *Model 2 - Demographic covariates added* |  |  |  |  |
|  | **MM Score** | **Trails B time** | **DV Time** | **PC1** |
| **MM Score** |  |  |  | 0,370 |
| **Trails B time** | 0,392 |  |  | 0,686 |
| **DV Time** | 0,209 | 0,420 |  | 0,586 |
| **DV Errors** | 0,116 | 0,108 | 0,076 | 0,222 |
|  |  |  |  | **R2=43,93%** |
| *Model 3 - Health covariates added* |  |  |  |  |
|  | **MM Score** | **Trails B time** | **DV Time** | **PC1** |
| **MM Score** |  |  |  | 0,371 |
| **Trails B time** | 0,367 |  |  | 0,687 |
| **DV Time** | 0,186 | 0,393 |  | 0,588 |
| **DV Errors** | 0,102 | 0,095 | 0,067 | 0,212 |
|  |  |  |  | **R2=42,65%** |
| *Model 4 - Quality of life covariates added* |  |  |  |  |
|  | **MM Score** | **Trails B time** | **DV Time** | **PC1** |
| **MM Score** |  |  |  | 0,367 |
| **Trails B time** | 0,357 |  |  | 0,688 |
| **DV Time** | 0,181 | 0,389 |  | 0,587 |
| **DV Errors** | 0,105 | 0,097 | 0,068 | 0,219 |
|  |  |  |  | **R2=42,45%** |

**Supplementary table S1**. Cognitive test intercorrelations, loadings on the first principal component, and variance accounted for this first principal component in the four models. Note that Mini-Mental scores have been multiplied by -1 so that higher values mean lower performance, as with the other tests.

|  | **No covariates** | **Demographics** | **+Health** |
| --- | --- | --- | --- |
| **Demographics** | 0.9002 |  |  |
| **+Health** | 0.8488 | 0.9432 |  |
| **+Quality of life** | 0.8415 | 0.9344 | 0.9910 |

**Supplementary table S2**. Intercorrelations of first principal component scores across the four models.

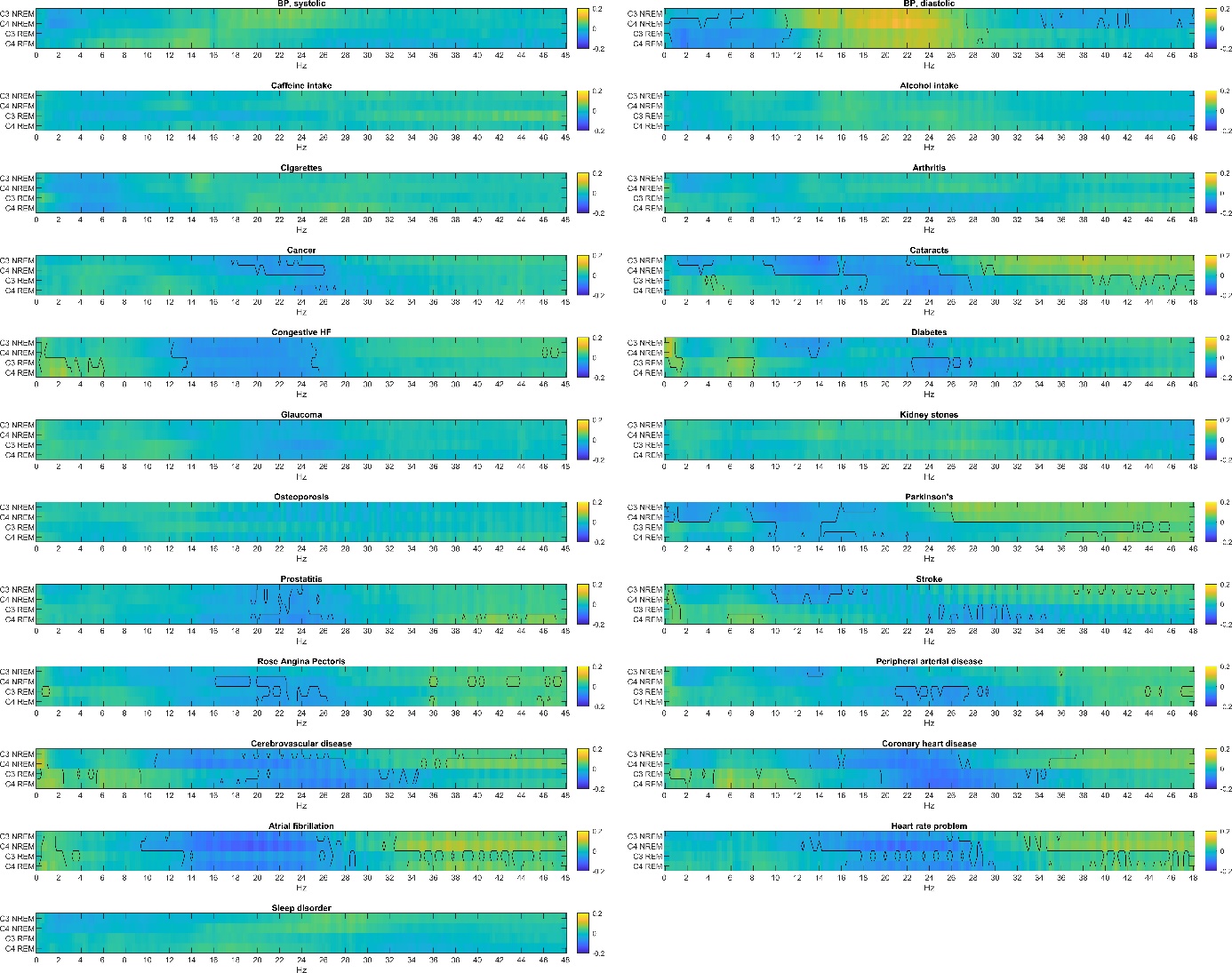

**Supplementary figure S1**. Point-biserial correlations between self-reported conditions and PSD. Black lines highlight correlations which are statistically significant after Benjamini-Hochberg correction for multiple comparisons.

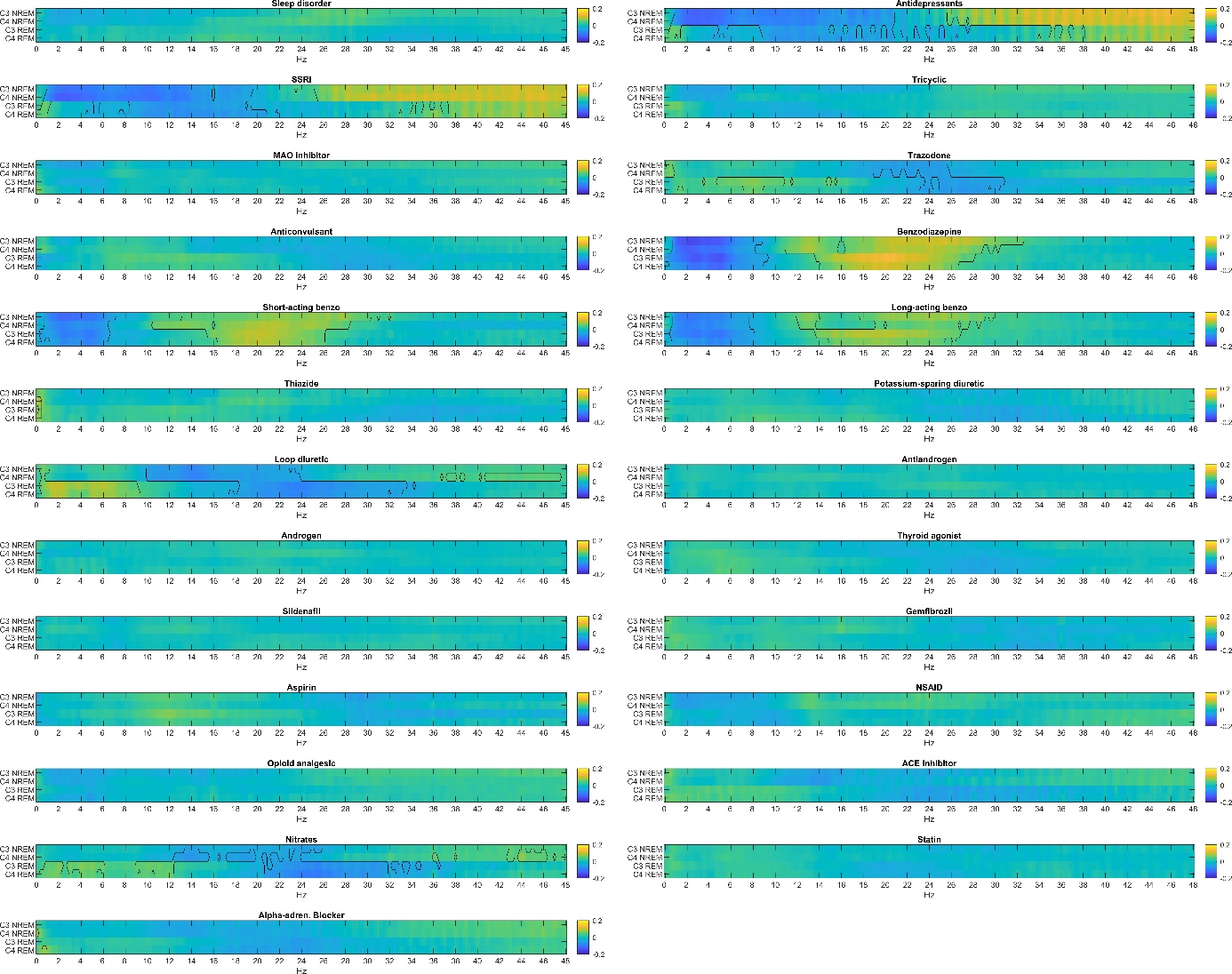

**Supplementary figure S2**. Point-biserial correlations between medication intake and PSD. Black lines highlight correlations which are statistically significant after Benjamini-Hochberg correction for multiple comparisons.

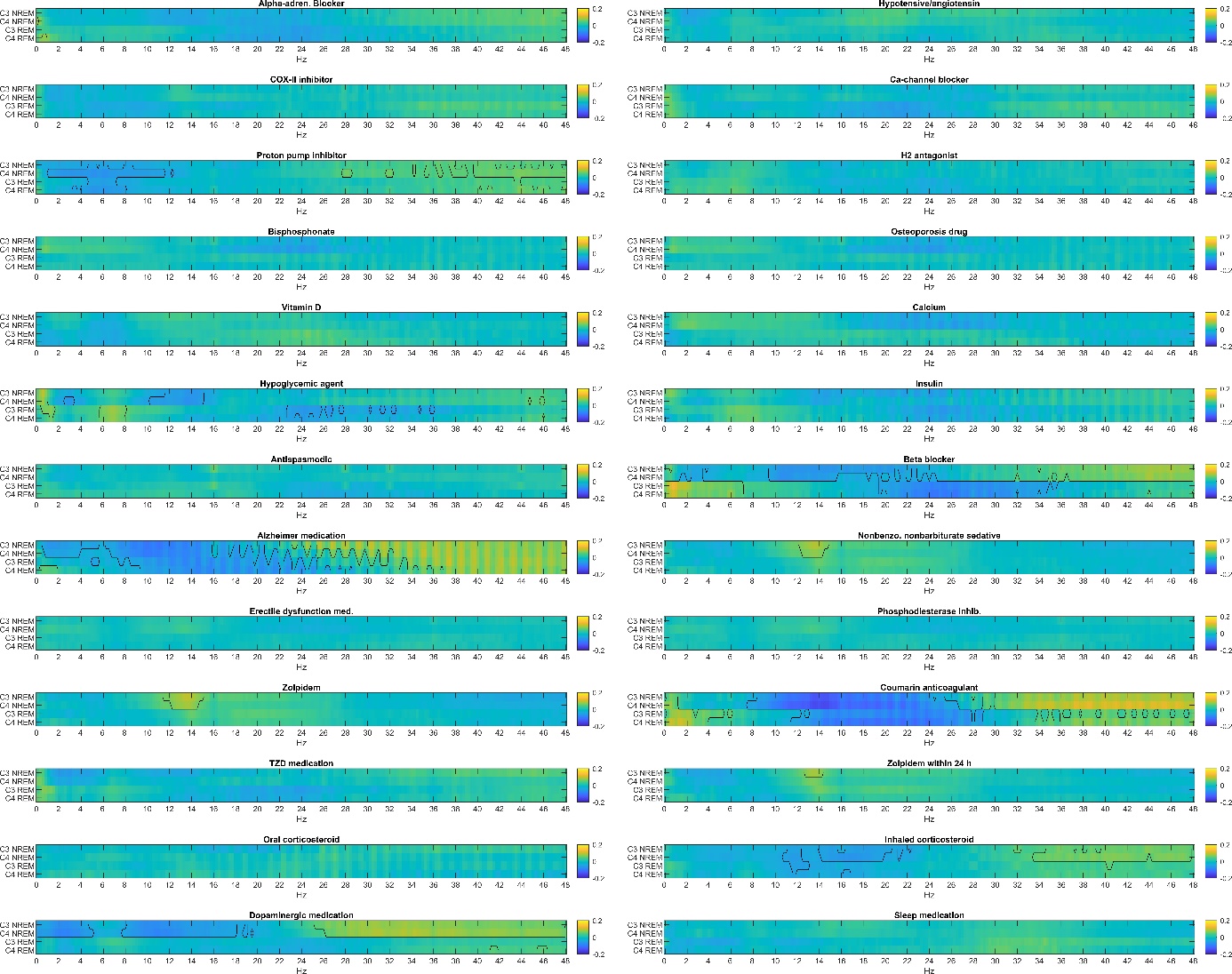

**Supplementary figure S3**. Point-biserial correlations between medication intake and PSD (continued). Black lines highlight correlations which are statistically significant after Benjamini-Hochberg correction for multiple comparisons.

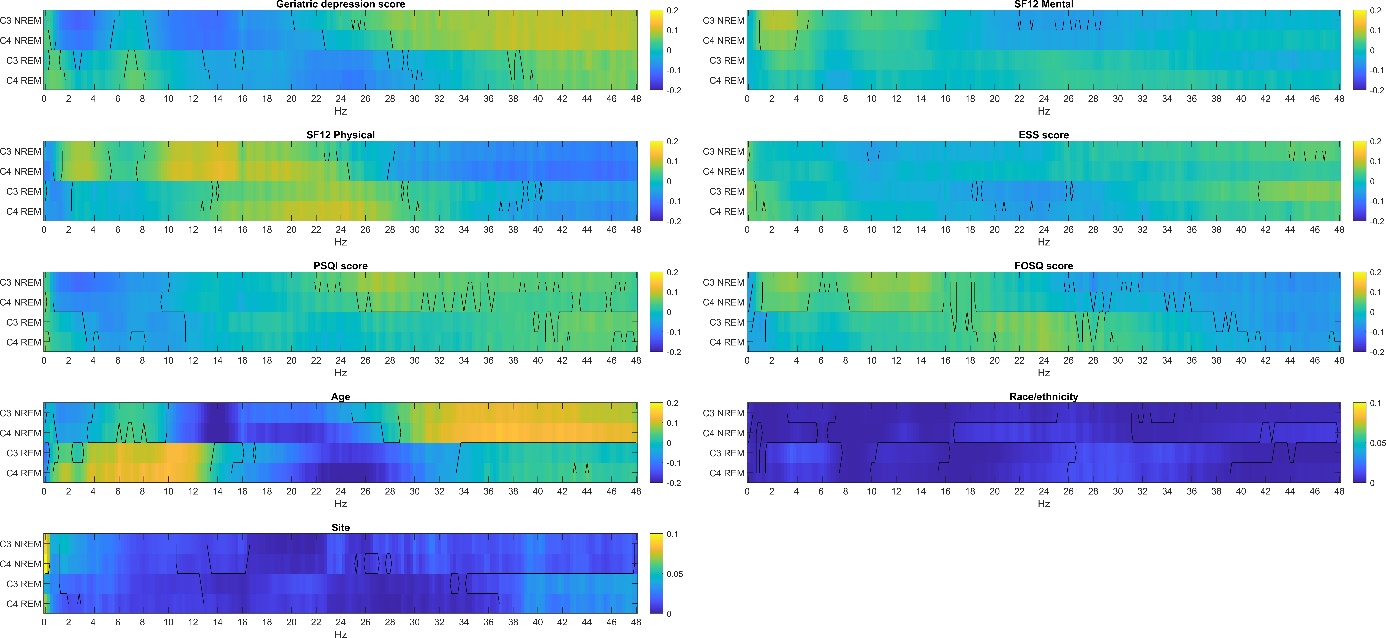

**Supplementary figure S4**. The association between quality of life variables, age, race/ethnicity and recording site. For categorical variables race/ethnicity and recording site intraclass correlations (PSD variance accounted for by category membership) are shown. For the other variables, Pearson correlations are shown. Black lines highlight correlations which are statistically significant after Benjamini-Hochberg correction for multiple comparisons.

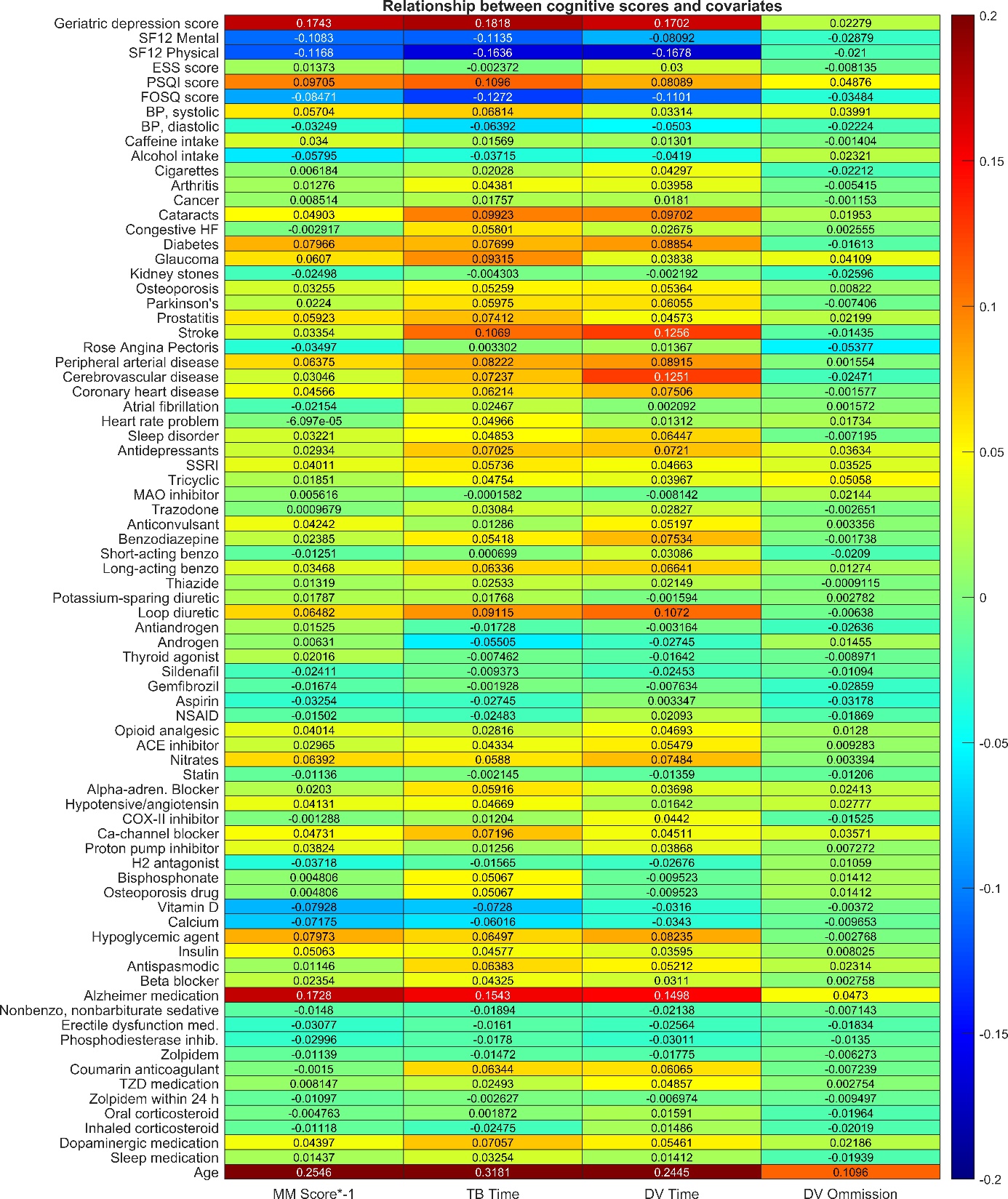

**Supplementary figure S5**. Correlations between covariates and cognitive test scores. 3MS scores have been multiplied by -1 to ensure that, as with the other scores, a positive correlation implies worse performance among those with a condition or with higher values of a continuous covariate. For the categorical covariates Race/ethnicity between-category variance was 4.4%, 2.4%, 0.6% and 0.9% for 3MS, Trails B completion time, Digit vigilance completion time and Digit vigilance completion errors, respectively. For categorical covariate Site, the same values were 0.8%, 0.4%, 1.6%, and 3.2%, respectively.
