## Supplementary text for "Multivariate prediction of cognitive performance from the sleep electroencephalogram"

**List of self-reported medications used as covariates:**

Any antidepressants, selective serotonine reuptake inhibitors, tricyclic antidepressants, monoamine oxidase inhibitors, trazodone, any anticonvulsants, any benzodiazepines, short-acting benzodiazepines, long-acting benzodiazepines, thiazide, potassium-sparing diuretics, loop diuretics, antiandrogens, androgens, thyroid agonists, sildenafil, gemfibrozil, aspirin, NSAID, opioid analgesics, ACE inhibitors, nitrates, statin, alpha-adrenergic blockers, hypotensive agents/angiotensin, COX-II inhibitors, calcium channel blockers, proton pump inhibitors, H2 antagonists, bisphosphonate, any osteoporosis drugs, vitamine D, calcium, hypoglycemic agents, insulin, uninary antispasmodics, beta blockers, Alzheimer’s disease medications, nonbenzo-nonbarbiturate sedative hypnotics, erectile dysfunction medications, phosphodiesterase type 5 inhibitors, Zolpidem in general, Zolpidem within 24 hours of PSG, anticoagulants (coumarin derivatives), TZD medications, inhaled corticosteroids, oral corticosteroids, dopaminergics, any sleep medications
